# Full-Length Envelope Analyzer (FLEA): A tool for longitudinal analysis of viral amplicons

**DOI:** 10.1101/230474

**Authors:** Kemal Eren, Steven Weaver, Robert Ketteringham, Morné Valentyn, Melissa Laird Smith, Venkatesh Kumar, Sanjay Mohan, Sergei L Kosakovsky Pond, Ben Murrell

## Abstract

Next generation sequencing of viral populations has advanced our understanding of viral population dynamics, the development of drug resistance, and escape from host immune responses. Many applications require complete gene sequences, which can be impossible to reconstruct from short reads. HIV-1 *env*, the protein of interest for HIV vaccine studies, is exceptionally challenging for long-read sequencing and analysis due to its length, high substitution rate, and extensive indel variation. While long-read sequencing is attractive in this setting, the analysis of such data is not well handled by existing methods. To address this, we introduce `FLEA` (Full-Length Envelope Analyzer), which performs end-to-end analysis and visualization of long-read sequencing data.

`FLEA` consists of both a pipeline (optionally run on a high-performance cluster), and a client-side web application that provides interactive results. The pipeline transforms FASTQ reads into high-quality consensus sequences (HQCSs) and uses them to build a codon-aware multiple sequence alignment. The resulting alignment is then used to infer phylogenies, selection pressure, and evolutionary dynamics. The web application provides publication-quality plots and interactive visualizations, including an annotated viral alignment browser, time series plots of evolutionary dynamics, visualizations of gene-wide selective pressures (such as *dN* /*dS*) across time and across protein structure, and a phylogenetic tree browser.

We demonstrate how `FLEA` may be used to process Pacific Biosciences HIV-1 *env* data and describe recent examples of its use. Simulations show how FLEA dramatically reduces the error rate of this sequencing platform, providing an accurate portrait of complex and variable HIV-1 *env* populations.

A public instance of `FLEA` is hosted at http://flea.datamonkey.org. The Python source code for the FLEA pipeline can be found at https://github.com/veg/flea-pipeline. The client-side application is available at https://github.com/veg/flea-web-app. A live demo of the P018 results can be found at http://flea.murrell.group/view/P018.

## Introduction

Next generation sequencing (NGS) has become an invaluable tool for studying HIV-1 and other rapidly evolving viruses by providing direct high resolution measurements of viral genetic diversity within the host. NGS has been used to study immune escape [1–7], drug resistance [7–13], transmission bottlenecks [10,14–16], population structure and dynamics [2,3,17–23], tropism dynamics [24], and multiplicity of infection [25]. It is also used in clinical virology [26,27]. For reviews of the promises and challenges of NGS applications in virology, see [28], [29], [30], and [31].

Full-length sequences can resolve features that are difficult to assemble from short sequences [8,32]. For instance, Pacific Biosciences SMRT sequences were able to resolve 1.5 kb *msg* isoforms from *Pneumocystis jirovecii*, but reads from a 454 instrument could not be assembled correctly [32]. For tracking evolutionary patterns in viral populations, accurately resolving these features provides a more accurate history of the population, which becomes especially relevant when epistatic interactions and linkage between mutations effect phenotypic changes in the pathogen [33–35]. For example, studies of HIV-1 *env* frequently use functional assays to measure the potency with which a given antibody or donor serum neutralizes a specific *env* strain [36], which requires knowing the full *env* sequence.

We have developed a pipeline for handling long read HIV-1 *env* sequencing data from within-host viral populations: the Full-Length Envelope Analyzer (`FLEA`). `FLEA` addresses the specific challenges posed by large volumes of such data, e.g., using the sequencing protocols we previously described in Laird *et al* [37], which also contains an overview of a prototype of `FLEA`. Here we describe the full pipeline and experimentally demonstrate its ability to resolve populations of closely related variants. `FLEA` uses state-of-the-art tools and methods at every step and can be accessed through a web browser or on a high-performance cluster. `FLEA` is readily extensible to other genes and systems.

`FLEA` has recently been used by the authors in two high-profile studies. In [38], we describe how `FLEA` was used to process PacBio HIV-1 *env* data from a clinical trial of monoclonal antibody 10-1074. For sequences sampled before and after therapy, `FLEA` reveals that prior to antibody therapy low-frequency *env* variants were present with mutations that typically confer resistance to 10-1074. Additionally, when resistance emerges, it emerges multiple times, exploiting many different resistance pathways. `FLEA` was also used to characterize the longitudinal *env* population that drove development of a broadly neutralizing antibodies against the apex of the *env* trimer, sampled from donor PC64 from the Protocol C primary infection cohort [39].

There exist dozens of standalone pipelines developed for analyzing HIV-1 and related sequence data, including longitudinal samples [4,9,13,40]. However, it was necessary to develop a new tool due to HIV-1 *env′s* extensive natural indel variation and the high rate of indels in long PacBio reads, which are especially problematic when any spurious indel in the 2.6kb *env* amplicon corrupts the reading frame, rendering the sequence uninterpretable. With HIV-1 *env*, the common strategy of mapping reads to a reference fails because the diversity in variable regions of *env*, predominantly driven by extensive indel processes, means that these regions in sampled reads lack homology to those in any heterologous reference sequence. Instead, `FLEA` relies on a fine-grained cluster-and-consensus strategy to remove spurious indels from reads. The task is related to Liang *et al.* (2016), but, rather than distinguishing a small number of variants at 81-91% identity, we must distinguish potentially hundreds of variants that differ by only a handful of bases.

In addition to a standalone application, `FLEA` is also available as an online resource that provides interactive visualizations for all its analyses. To allow researchers to further examine and dissect their results, `FLEA` also provides access to raw data, such as aligned consensus sequences and phylogenetic trees.

## Design and Implementation

### Pipeline

The input to `FLEA` is a set of FASTQ files from the PacBio RS-II or Sequel. Each set corresponds to one time point, containing circular consensus sequence (CCS) reads, which can be obtained using the “Reads of Insert” protocol on PacBio’s SMRTportal or SMRTanalysis tools. Upon completion, the `FLEA` pipeline produces results as JSON (Javascript Object Notation) files, a standard format for machine (and human-) readable structured data. The logic of `FLEA` is implemented in `Nextflow` [41], a workflow framework for deploying parallel pipelines to clusters and clouds.

`FLEA` consists of multiple sub-pipelines, as shown in Fig. 1. Details of the quality and consensus pipelines are depicted in Fig. 2. Together, these two pipelines take error-prone CCS reads and convert them into unique high-quality consensus sequences. The alignment pipeline generates a multiple sequence alignment, which is used by multiple methods in the analysis pipeline.

**Figure 1.**
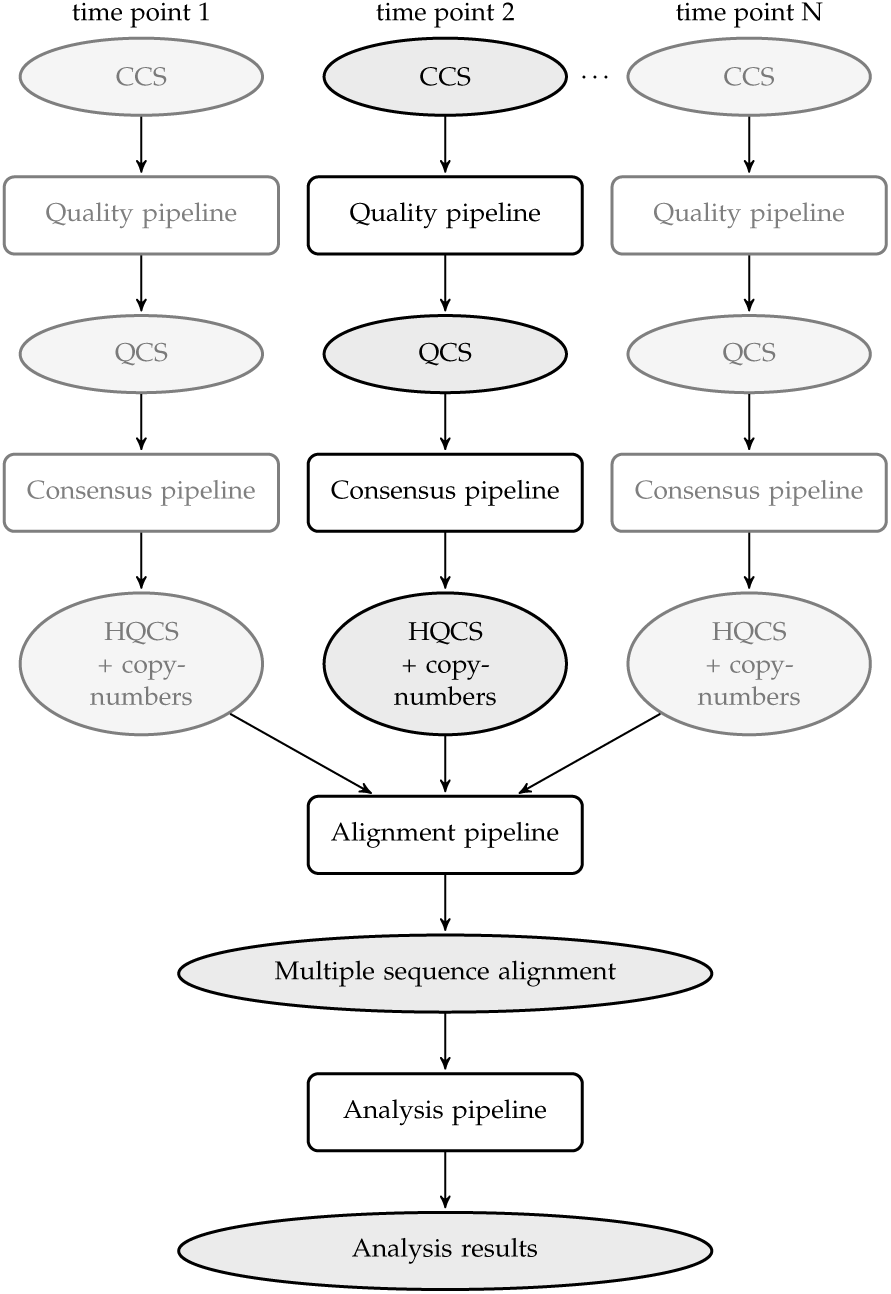
*Overview of the entire pipeline, broken into conceptual sub-pipelines. The* Quality *and* Consensus *subpipelines process each time point separately. Duplicate steps in other time points are grayed out. CCS stands for “circular consensus sequences”; QCS for “quality-controlled sequences”, and HQCS for “high-quality consensus sequences”.*

**Figure 2.**
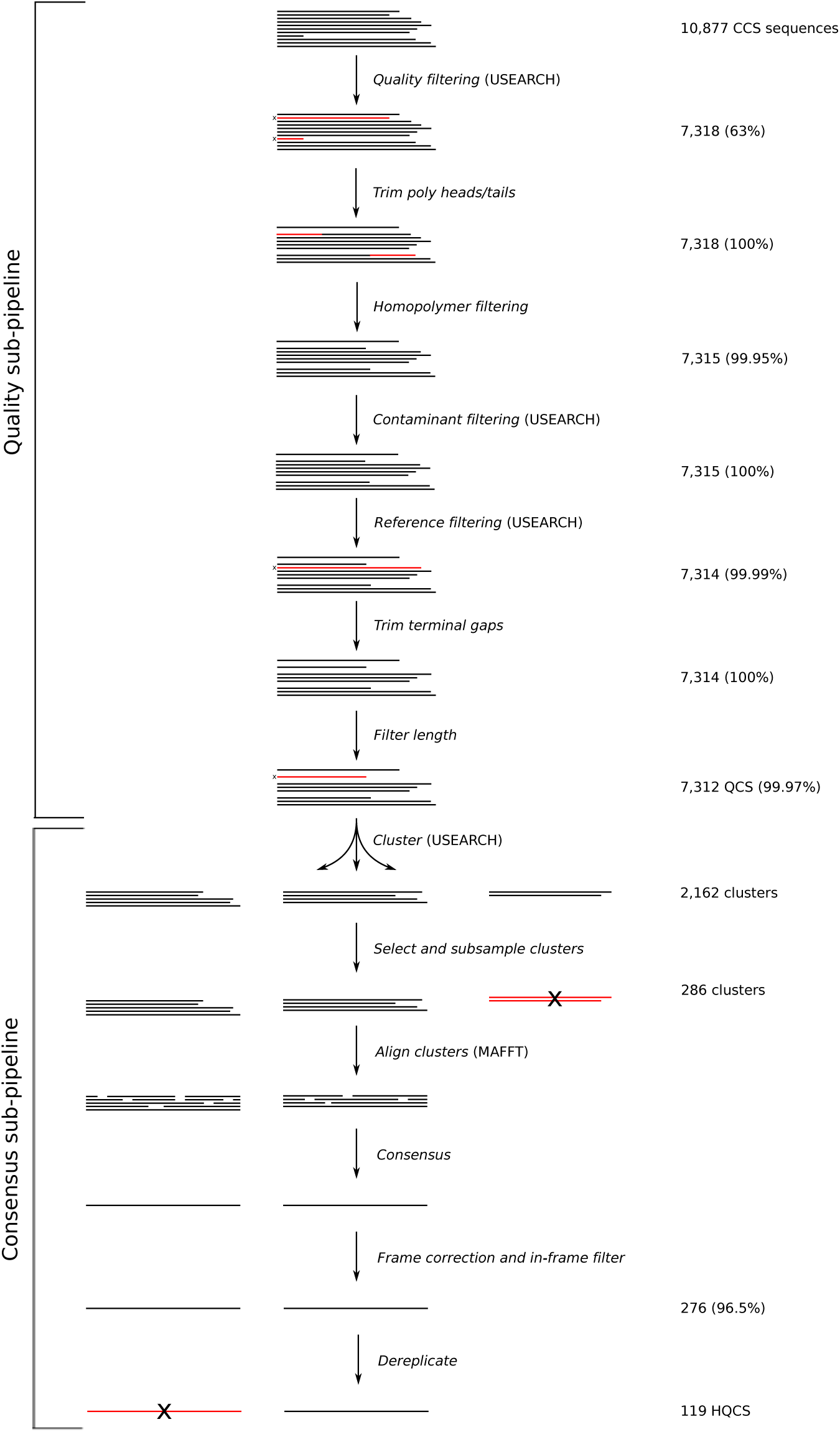
*Combined view of the quality and consensus sub-pipelines. These steps are repeated independently on each time point. Numbers are reported from the analysis of sequences from the first time point (V03) of donor P018, which is 3 months post infection. Percentages give the fraction of sequences retained after filtering. Tasks indicate whether they use third-party tools **`USEARCH`** or **`MAFFT`***.

### Quality assurance sub-pipeline

The first steps remove low quality reads and filter out common sequencing artifacts. Parameters given in these steps were chosen for full-length HIV-1 envelope sequences from the RS-II or Sequel platforms. Other reads with different properties (error rates, error models, lengths, homopolymer distributions, etc.) likely require different parameters. All steps are run independently per time point.

1. **Filter by error rate.** The input FASTQ files contain Phred scores for each base, encoding the probabilities of incorrect base calls. USEARCH [42] is used to remove reads with an expected error rate greater than 1%, computed as the mean of the per-base error probabilities.
2. **Trim heads/tails.** A fraction of reads from the Laird *et al.* sequencing protocol contain poly-A or poly-T heads or tails (cause unknown), which can be hundreds of bases long and sometimes contain a small number of other bases. These heads and tails are trimmed with a hidden Markov model (Fig. 3) implemented in Pomegranate [43]. The emission probabilities of the model were fixed, and the transitions trained using Baum-Welch. The Viterbi path for each sequence is computed, and bases emitted by head and tail nodes are removed.
3. **Filter long runs.** Reads with homonucleotide runs longer than 16 bases are discarded. This length was chosen to be twice the length of the longest such run in the LANL HIV database [44].
4. **Filter contaminants and trim reads.** Sample contamination can introduce non-native sequences that interfere with subsequent analyses, so these contaminants must identified and discarded. `USEARCH` is used to compare reads to a contaminant database and a reference database using `usearch_global`. Alignments returned from querying the database are then used to trim reads to the gene boundaries. Trimming terminal insertions is vital for the accuracy of downstream tasks, such as length filtering and clustering. The contaminant database contains HXB2 and NL4-3 *env*, each ubiquitous in labs working with *env* sequences and a common source of sample contamination. Reads that match with ≥ 98% identity are discarded. Since a 1% error rate cutoff was earlier used, this parameter conservatively ensures that these contaminants are almost certainly identified. The reference database contains thirty-eight sequences representing the major HIV-1 Group M subtypes from the LANL sequence database [44]. Reads with ≤ 70% identity to every sequence in the reference database are discarded. This cutoff is chosen to retain reads remotely similar to HIV-1 Group M while excluding contaminants such as human or bacterial genome reads. If a sample is from SIV, or from a non group-M HIV+ donor, then more appropriate reference sequences should be added to the database.
5. **Filter by length.** By default, sequences shorter than 90% or longer than 110% of the length of the reference sequence are discarded. However, sequences with large deletions are frequently observed in HIV. These likely represent replication incompetent envelopes, and their reduced length can cause them to be dramatically oversampled due to PCR length bias. Users who want to include these species in their analyses should modify these parameters.

**Figure 3.**
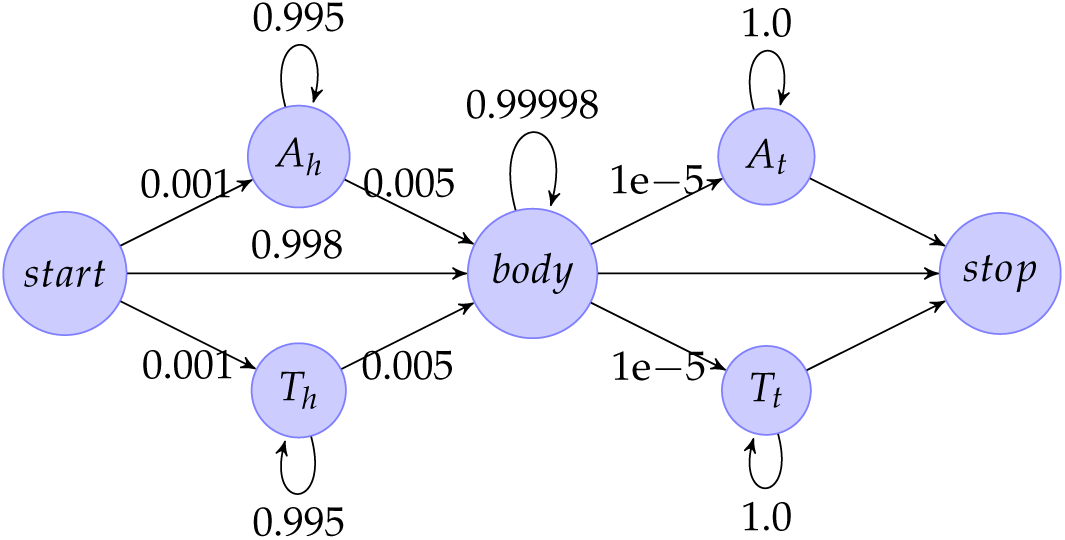
*Hidden Markov model used for trimming poly-A and poly-T heads and tails. A head and tail states have a small (p = 0.01) probability to emit non-A bases, and similarly for T. The body state emits all four bases with equal probability. The start, and stop states emit nothing.*

Reads that pass this quality assurance phase have low expected error rates and no homonucleotide runs, are within 70% identity of at least one reference sequence, are (after trimming) no more than 10% different in length than a reference sequence, and do not match the contaminant database. We refer to these sequences as quality-controlled sequences (QCS).

### Consensus sub-pipeline for variant identification

Even for highly diverse populations, unique reads in a sequencing run outnumber the true unique variants, predominantly due to sequencing errors. The problem is far more significant in long reads than in short reads, precluding the use of amplicon denoising strategies used to reduce error rates in short read sequencing [45]. To accommodate this effect, the next phase of the `FLEA` pipeline clusters and combines QCS reads, attempting to infer the true variants in each time point. It also attempts to detect and correct frameshift errors.

1. **Cluster.** `USEARCH` is used with the `cluster_fast` command to generate clusters with 99% nucleotide identity. This parameter approximates the 1% error cutoff used in the error rate filtering step, so that pairwise distances of sequences in the same cluster are consistent with the sequencing error. `cluster_fast` runs in a single pass, so it is sensitive to input order. Sequences are sorted from lowest to highest quality according to expected error rate; experiment suggests that this order yields better results (see supporting information).
2. **Select and subsample clusters.** Clusters with fewer than three members are discarded, because they are too small to de-noise by majority consensus. Clusters with more than 50 members are subsampled to the top 50 with the lowest expected error rate to speed up the multiple sequence alignment step.
3. **Align and consensus.** `MAFFT` [46] is used to align each cluster. The consensus sequence of each alignment is computed.
4. **Frame correction** In-frame consensus sequences from all time points are collected into a USEARCH database for frame correction. `usearch_global` is then used to align each out-of-frame sequence to its top hit. The nucleotide alignment is used to correct incomplete codons: short insertions (1 or 2 base pairs) are discarded, and single deletions are replaced with the aligned base. Sequences with longer insertions or deletions are discarded. All changes are logged, so that the user can identify the sequences that have been corrected.
5. **Uniqueness** Non-unique consensus sequences are dereplicated using `usearch-fastx_uniques`.
6. **Copy numbers** The number of sequences per cluster provides an estimate of the relative abundance of that HQCS in the population. Those numbers are further augmented by adding sequences orphaned by cluster filtering and HQCS dereplication. `usearch_global` is used to assign each QCS to its nearest HQCS. The number of sequences accrued by each HQCS is interpreted as its copy number.

All of these tasks are run separately for each time point, yielding sets of unique in-frame consensus sequences. We refer to these sequences as high-quality consensus sequences (HQCS).

### Alignment sub-pipeline

The HQCSs from all time points are combined into a single file, translated to protein sequences, and aligned using `MAFFT`. A Python script then transfers the gaps from each aligned protein sequence to the corresponding nucleotide sequences to produce a codon-level nucleotide multiple sequence alignment of all unique variants from all time points.

### Analysis sub-pipeline

The analyses used in `FLEA` take as input the two outputs of the alignment phase: a codon multiple sequence alignment of all unique HQCS sequences from all time points, and their associated copy numbers. These data are used for the following analyses.

1. **Time point metrics.** `HyPhy` [47] scripts are used to compute evolutionary metrics (total, *dN*, and *dS* divergence and diversity) and phenotypic metrics (protein length, potential N-linked glycosylation sites, isoelectric point) for each annotated region (e.g., V1, MPER) in the amplicon for each time point.
2. **MRCA.** The most recent common ancestor is inferred by taking the copy-number-weighted codon consensus of the codon-aligned HQCSs from the earliest time point. By including gaps, the MRCA sequence is already aligned with the rest of the multiple sequence alignment. This strategy is acceptable for primary infection studies from single founders with very low early diversity.
3. **Reference coordinates.** `MAFFT` is used to assign HXB2 [48] coordinates to the gapped MRCA sequence, which are then transferred to the full multiple sequence alignment.
4. **Infer phylogeny.** A maximum-likelihood phylogenetic tree is inferred with FastTree2 [49,50] under the general time reversible model.
5. **Ancestral sequence reconstruction.** `HyPhy` is used to infer ancestral sequences at the internal nodes of the phylogeny, using joint maximum likelihood reconstruction and the HKY85 substitution model [51].
6. **Multidimensional scaling.** TN93 [52] is used to compute a distance matrix for all HCQC sequences using the Tamura Nei 93 distance [53]. Metric multidimensional scaling [54] (implemented in `scikit-learn` [55]) is used to find a two-dimensional embedding of the sequences that approximates their pairwise distances.
7. **FUBAR.** Site-specific selection rates are inferred using `FUBAR` [56], implemented in `HyPhy`.
8. **Position-specific changes.** Entropy and Jensen-Shannon divergence are computed for each position in each time point.

The results of these analyses are provided to the user in an interactive web application, described next.

## Web application

The `FLEA` web app is built using modern web design principles. It consists of two parts: a Javascript client-side app, written using the `Ember.js` [57] framework, and a server-side REST (REpresentational State Transfer) service for serving JSON-formatted data. There are two main benefits to using this decoupled pattern for scientific web applications. First, the client-side code only needs to be downloaded once, at the start of the session. The data are requested from the server and cached as needed. Once everything is loaded, the visualizations run entirely in the browser with no delays for page loads. Second, the REST service may be reused by other apps and third-party tools.

The web app presents the results of the `FLEA` analysis as a series of interactive visualizations. The report is organized into the following sections.

### Multidimensional scaling

A two dimensional embedding of the HQCSs is visualized as a bubble plot, showing changes in population structure over time, as shown in Fig. 4. This visualization has been especially useful for investigating populations with superinfection, or with multiple founders, where aggressive recombination between vastly different *env* variants precludes the use of phylogenies.

**Figure 4.**
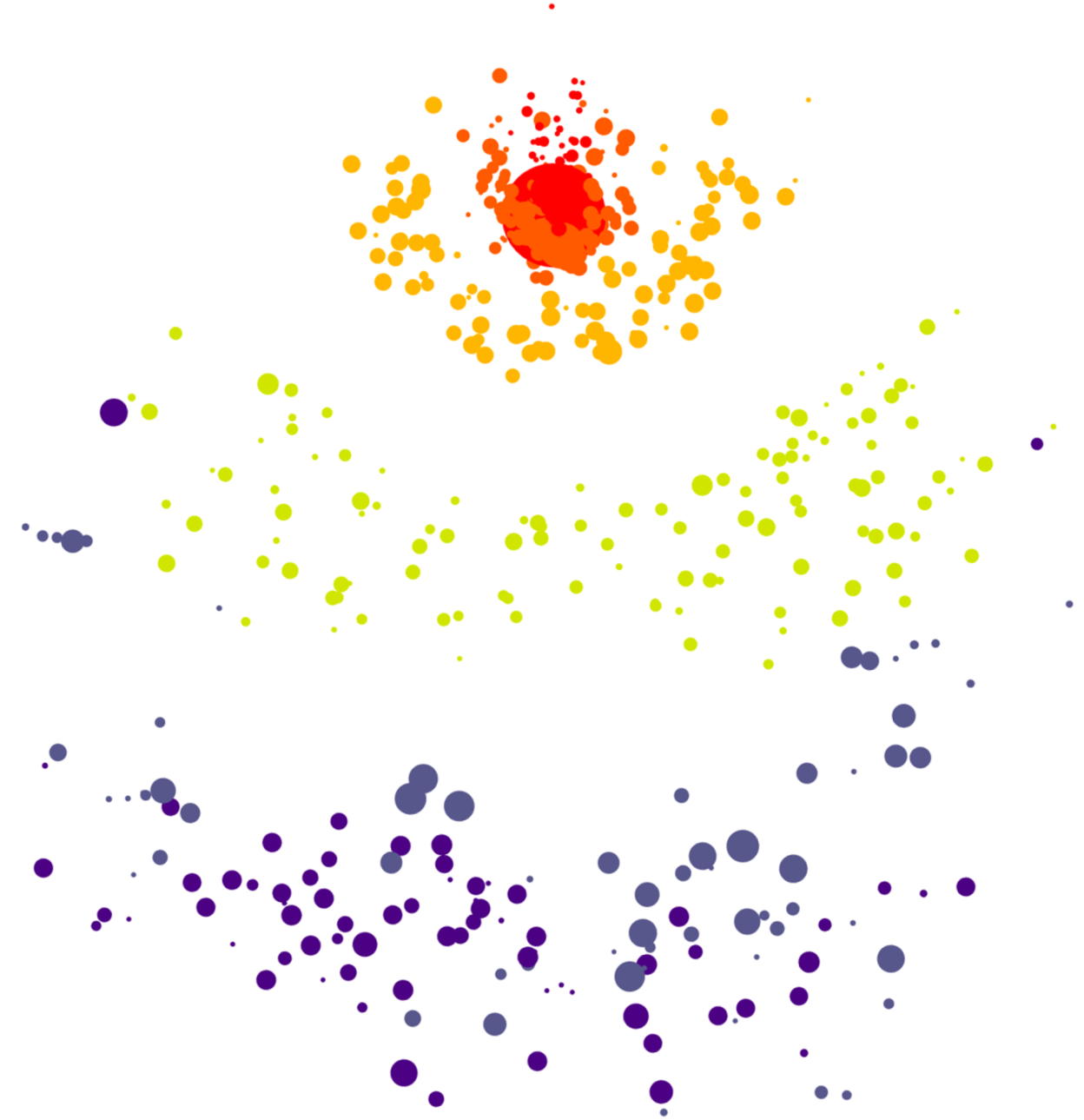
*Screenshot of the multidimensional scaling plot. The embedding in two dimensions preserves pairwise evolutionary distances between HQCSs. Node area is proportional to copy number, and color corresponds to time point. The increasing genetic diversity of the population is visible as time goes on.*

### Evolutionary trajectory

The evolutionary trajectory viewer plots evolutionary and phenotypic metrics for each time point and multiple regions in the amplicon, giving a high-level overview of population dynamics over time. Fig. 5 shows the plot for the entire gp160 region of HIV-1 Env, which is generated with the `D3.js` plotting library [58].

**Figure 5.**
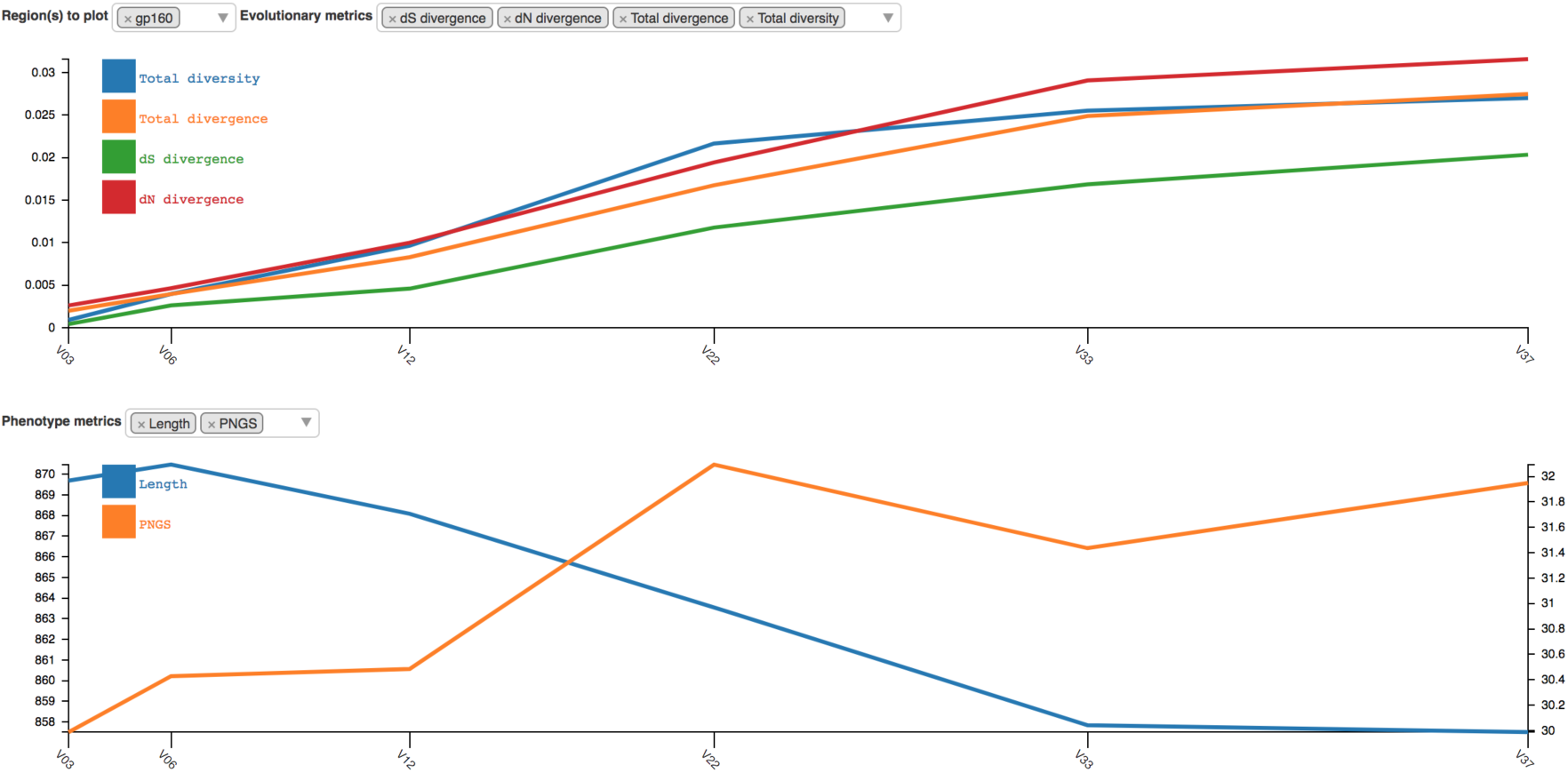
*Screenshot of the evolutionary trajectory report. Four evolutionary metrics (dS divergence, dN divergence, total divergence, and total diversity) and two phenotype metrics (length and possible N-linked glycosylation sites) are shown for gp160.*

### Sequences

The multiple sequence alignment of all the HQCSs sequences is the foundation for all subsequent analyses. It is displayed in the amino acid sequences viewer, which contains a custom alignment browser and an interactive motif dynamics plot, as shown in Fig. 6.

**Figure 6.**
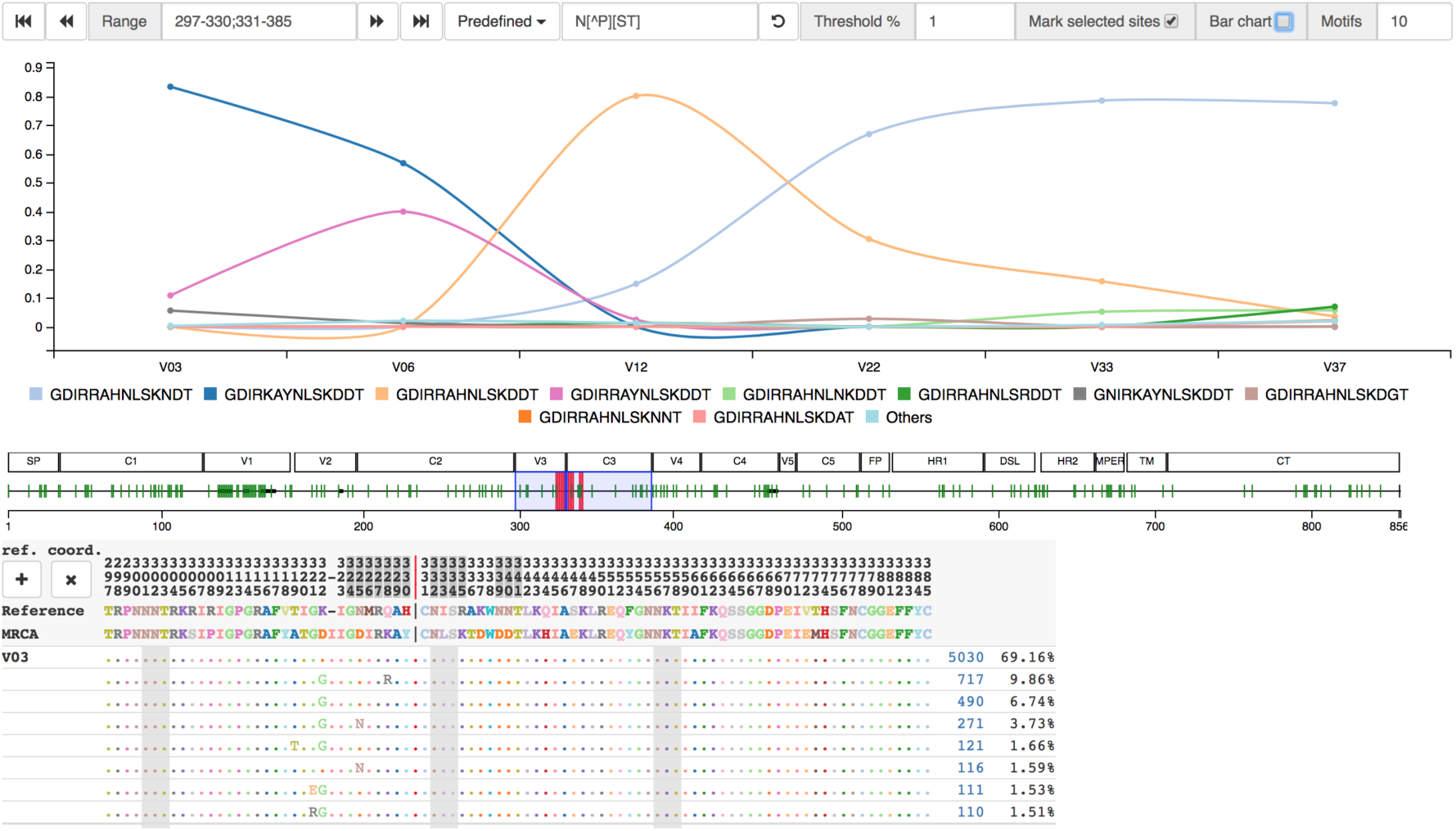
*Screenshot of amino acid sequences viewer. Sequences are grouped by identity, with aggregate copy number and population percentage shown to the right. An overview of the amplicon, optionally annotated with region names, provides fast access to different locations of the alignment. Selecting columns of the alignment interactively updates the amino acid dynamics plot, showing the dynamics of the selected motif over time. In this case, the trajectory shows changes in the N332 glycan supersite. Sites inferred by FUBAR to be undergoing positive selection are selectable*.

### Protein structure

The protein structure viewer maps evolutionary metrics to an interactive three-dimensional structure of the protein, customized from PDB ID `5FUU`, a recently resolved cryo-EM structure [59], and rendered using `pv` [60]. Missing residues are rendered as spheres which are positioned by Bézier curve interpolation. *dN*/*dS* ratios, Jensen Shannon divergence, and entropy may all be mapped to the protein structure, as shown in Fig. 7. The same metrics are also plotted in one dimension for each time point, as shown in Fig. 8. The protein visualization interacts with the sequence viewer by showing alignment positions and highlighting the residues in the selected sequence motif.

**Figure 7.**
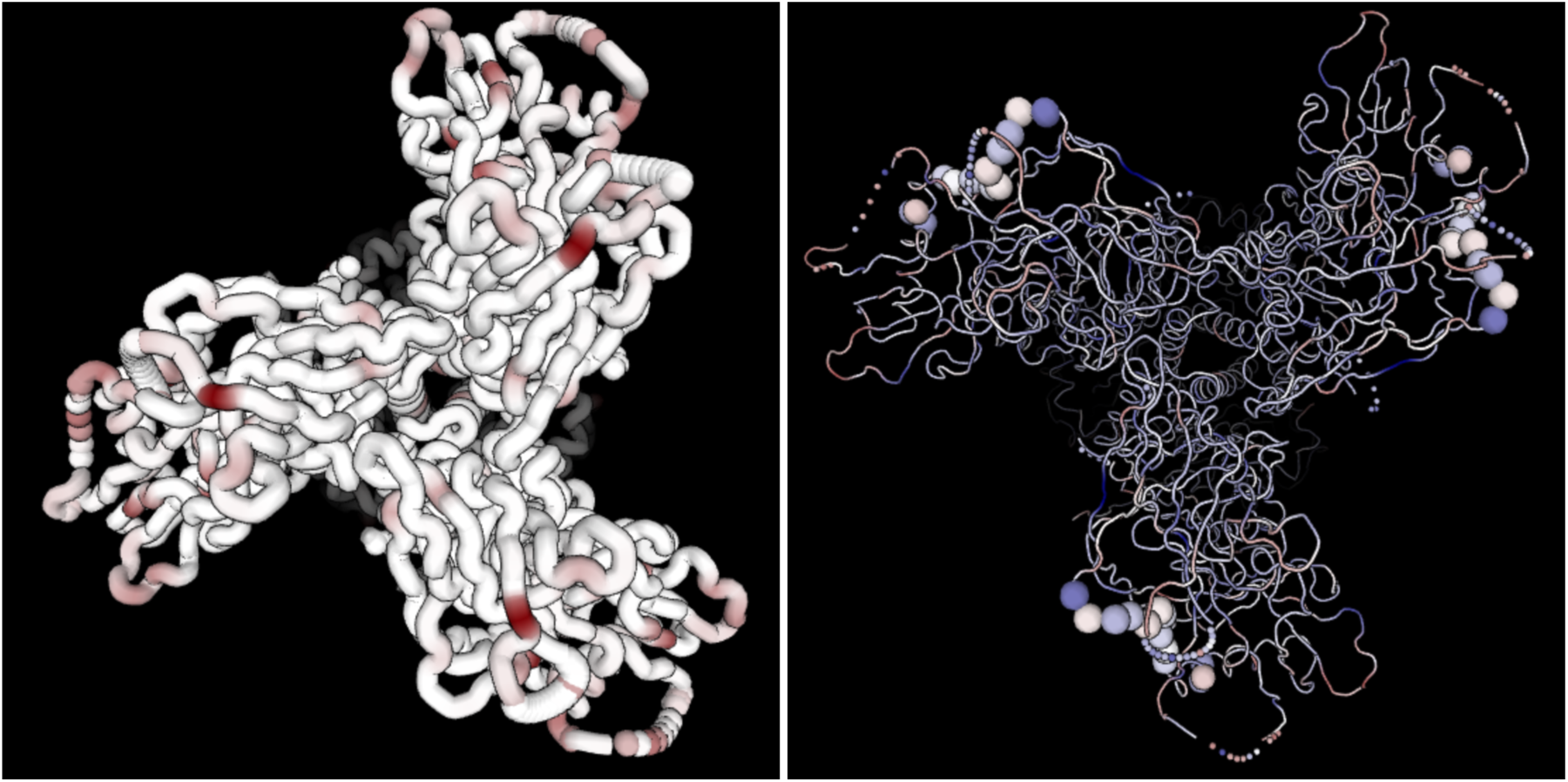
*Screenshots of the interactive three-dimensional Env structure, colored according to JS divergence (left) and dN/dS values (right). Positions imputed to be undergoing more positive selection (dN/dS > 1) are darker red, and positions undergoing more purifying selection (dN/dS < 1) are darker blue. The right structure also shows motif positions highlighted in the sequence viewer*.

**Figure 8.**
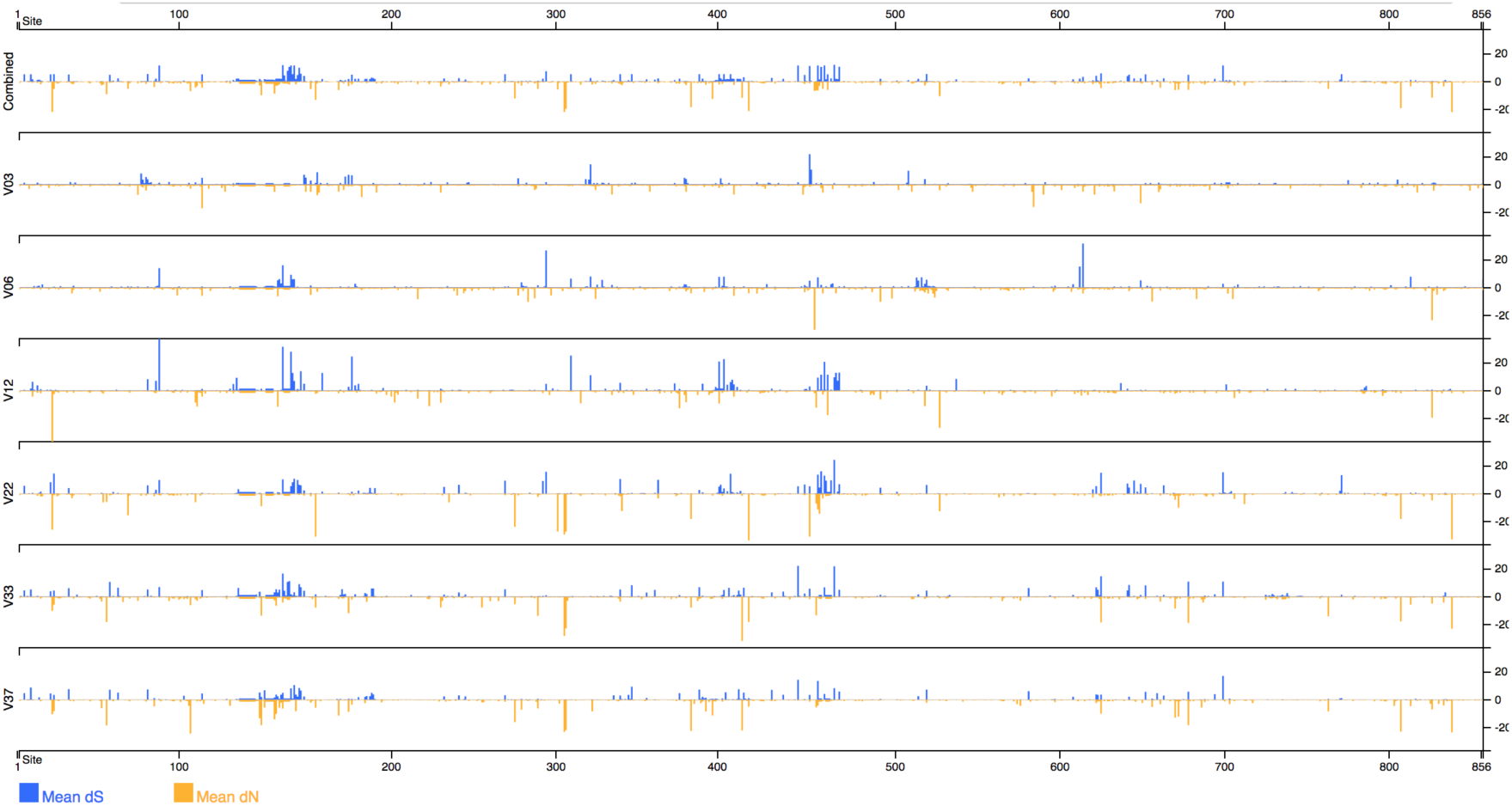
*Screenshot of dN/dS values mapped to protein positions and separated by time point*.

### Trees

The tree viewer renders a tree browser with `phylotree.js` [61], as shown in Fig. 9. Leaf nodes are scaled to the copy number of their sequence. The tree zoom level, layout, and coloring is interactively modifiable. Motifs selected in the sequence viewer are mapped to the tree. Ancestral nodes are colored by motif, allowing inferred changes to be tracked through the entire phylogeny.

**Figure 9.**
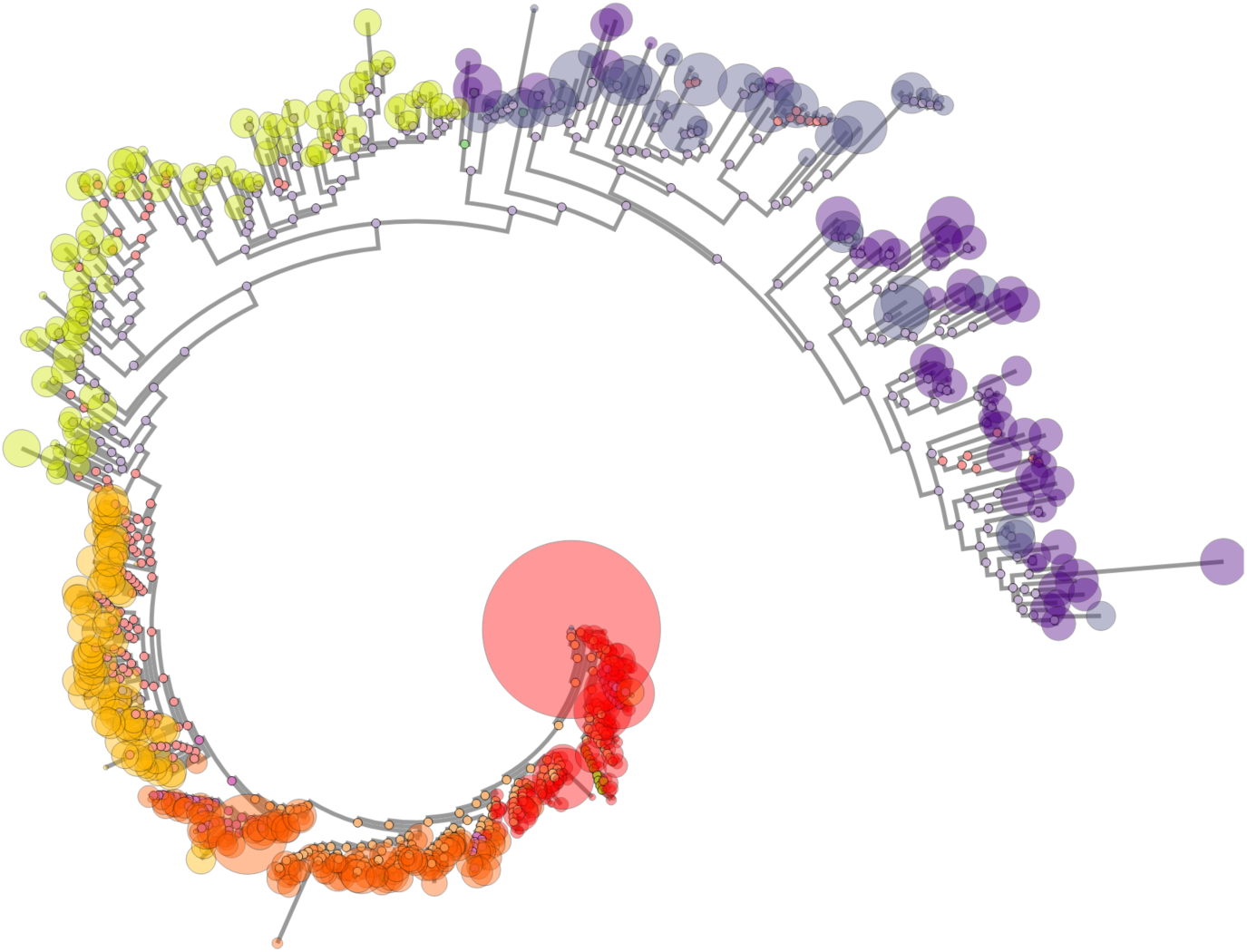
*Screenshot of the phylogenetic tree viewer. Leaf node size corresponds to sequence copy number. Node color corresponds to time point. Since ancestral sequences have been inferred, ancestral nodes are colored according to the selected motif, which in this case is the N332 glycan supersite*.

## Results

The entire pipeline was run on HIV-1 *env* reads from donor P018, which are available from the NCBI Sequence Read Archive under BioProject PRJNA320111, and were sequenced as part of [37] on the RS-II instrument, using the older generation P5/C3 PacBio sequencing chemistry. The full dataset contains 58,468 CCS reads. The reads are split across six time points, which are coded as V03, V06, V12, V22, V33, and V37, where *Vx* corresponds to a visit x months post infection. The number of reads per time point ranges from 7,530 in V33 to 11,806 in V06.

### Results on simulated data

The true sequences and copy numbers are not known for the P018 data. In order to assess the accuracy of our inferred sequence population, we used the HQCSs from a previous `FLEA` run to simulate a gold standard dataset on which to assess the `FLEA` pipeline.

The simulation procedure starts with the HQCSs and copy numbers from the `FLEA` results on P018, then augments them with additional mutated sequences to create a gold standard set of templates. Mutated sequences were added because our clustering strategy may artificially merge similar templates. For each template, noisy reads with a SMRT-style error profile were sampled. Full details of the simulation process appear in the supporting information. These simulated reads were sent through the `FLEA` pipeline, both with and without frame correction.

The resulting QCS and HQCS sequences were compared to the ground truth using Earth Mover’s Distance (EMD), using normalized copy numbers for the population weights and edit distance for the distance matrix. The fully constrained EMD has units that can be directly interpreted as the average change per nucleotide necessary to transform one sequence population into another. We also calculate two variants of EMD for further insight into how well the inferred population *B* estimates the sequences in the ground truth population *A. EMD_FP_* removes the constraint on *A*, allowing any amount of flow from *A* to *B*. It is a measure of false positives because it grows when *B* contains extra sequences distant from any in *A.* Similarly, *EMD_FN_*removes the constraint on *B*. It grows when *B* fails to recapitulate sequences in *A*, and therefore is a measure of false negatives.

To see the effect of sequencing runs of different depths, the experiment was repeated for 300,1, 000, 3,000, and 10,000 reads per time point. The results, which appear in Table 1, show the benefit of `FLEA`’s approach of reducing sequence errors via clustering and consensus. The QCS sequences, although they have few false negatives (*EMD_FN_* = 0.0782) for *n* = 10,000, are dominated by false positives (*EMD_FP_* = 8.3). However, adding the consensus sub-pipeline virtually eliminates false positives (*EMD_FP_* = 0.0336), at the cost of only a 2.4x increase in false negatives, for a 8.6x improvement in EMD to 1.0549. The frame correction step further improve both *EMD_FP_* and *EMD_FN_* because it turns false positives into true positives.

**Table 1.**
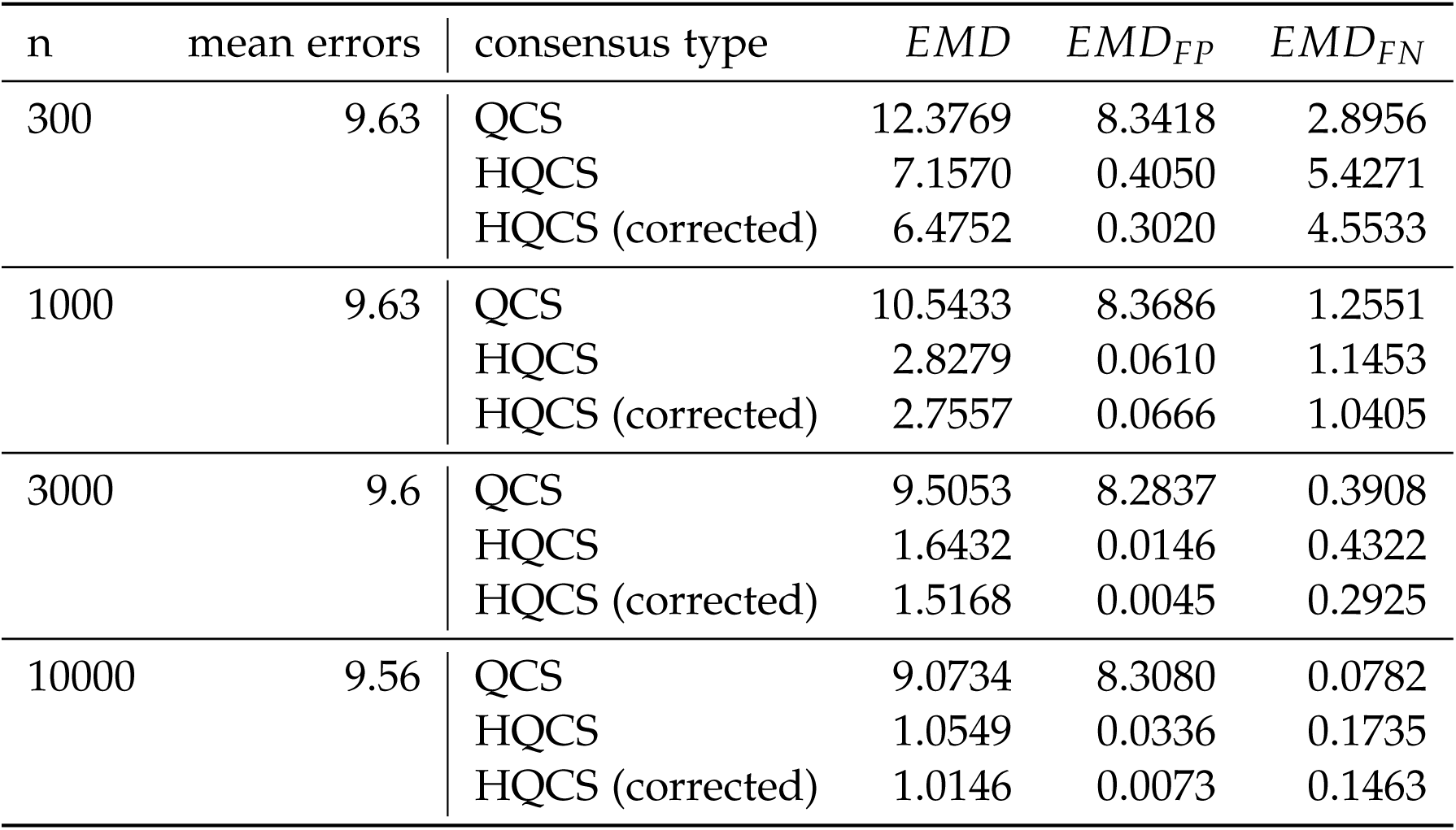
*EMD metrics for various numbers of reads, averaged across all time points. “mean errors” gives the average number of errors in the reads, estimated from the simulated Phred scores*.

The full-length *env* sequencing protocol yields approximately 10,000 reads per run; the P018 data averaged 9,744 reads per time point. Therefore, these results with *n* = 10,000 suggest that `FLEA` is capable of taking a full sequencing run of CCS reads from a diverse viral population with an average of 9.56 errors per sequence and inferring HQCSs with an average of 1.01 errors per sequence, which corresponds to an average error rate of 0.038%. Moreover, these error rates are mostly caused by low-abundance sequences in both the true population and the inferred `FLEA` sequences. Figure 10 shows that `FLEA` perfectly recovers all sequences from all time points that account for at least 1.6% of the population.

**Figure 10.**
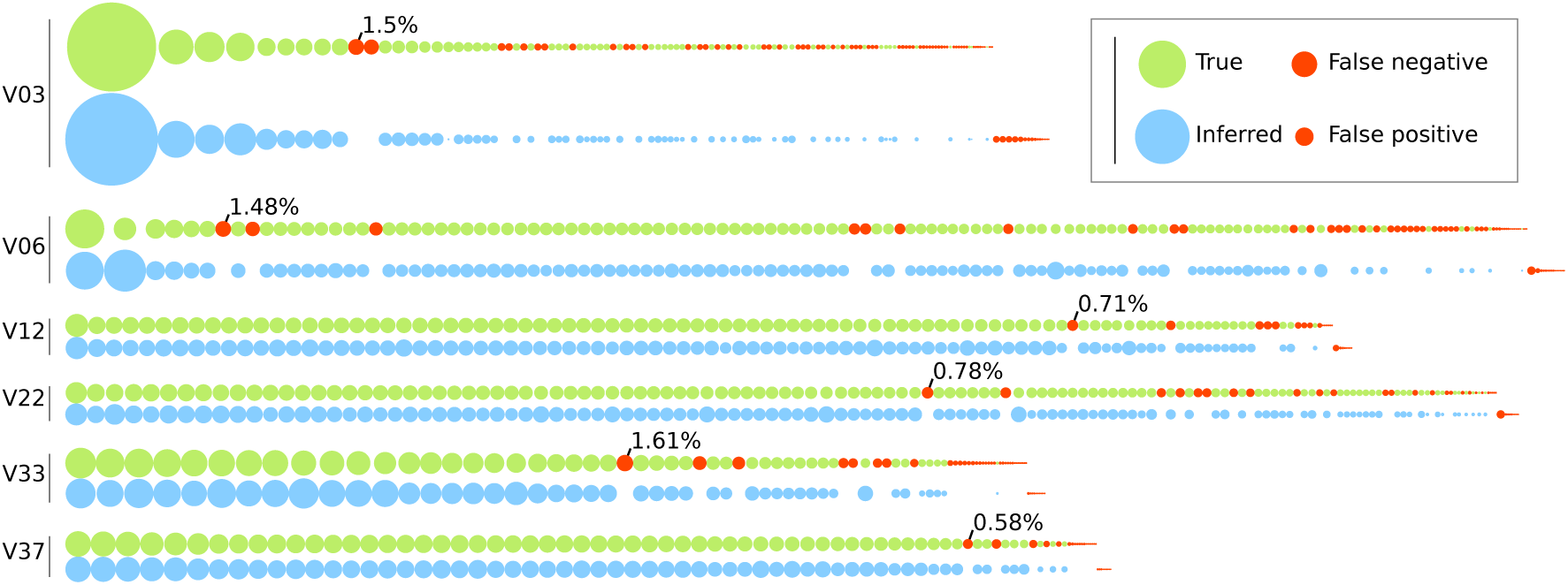
*Comparison of true sequence abundances versus copy numbers inferred by **`FLEA`** for each time point of the simulated P018 data. Each node represents one sequence, with the area denoting its relative abundance in the population. The true population (top) is colored green. For each true sequence, the matching HQCS sequences appears below it in blue. Red nodes denote false negatives and positives. The most common false negative for each time point is annotated with its abundance*.

### Results on real data: donor P018

`FLEA` was run directly on the P018 sequences, and the results are summarized here. The full results of this run are available to view at http://FLEA.murrell.group/view/P018..

Fig. 2 shows the number of sequences from the V03 time point that make it to each stage of the quality and consensus pipelines. At three months post infection, the majority amino-acid sequence variant is shared by 52.1% of the population, and the next most common variants accounts for just 8.66%. This relative lack of diversity is consistent with early infection dynamics. By 37 months post infection there is much more diversity: the most common variant accounts for only 3.96% of the population.

Donor P018 shows signs of potential N332 glycan specificity, as shown by the motif trajectories in Fig. 6. The glycan supersite, centered around N332 in V3, is a common target for broadly-neutralizing antibodies [62] because they are often conserved, so mutations in these regions are associated with escape [63]. A year into sampling (V12), mutations 328R and 330H dominate, and the majority of sequences also contain 339N from 22 months (V22) onwards.

## Discussion

The `FLEA` pipeline analyzes longitudinal full-length *env* sequences and provides visualizations of the results. Using simulations, we show that `FLEA` is capable of inferring accurate HIV-1 *env* consensus sequences and population frequencies. Despite each CCS read containing an average of ten errors, our approach distinguishes variants that differ by as little as one base from an amplicon with high indel variation. It uses those high-quality consensus sequences to generate a codon-aware multiple sequence alignment of all time points, estimate ancestral sequences, infer the phylogenetic tree, and perform many other population-level analyses with high accuracy. These results are presented in a visualization suite that is highly general and applicable to many related sequencing problem.

While our `USEARCH`-based clustering and consensus strategy for de-noising long PacBio amplicons performs well when error rates are < 1%, there is a clear need for more sophisticated long-read de-noising algorithms that exploit the additional depth of lower quality reads that we currently discard. This will be especially beneficial for longer PacBio amplicons, because the CCS read quality distribution degrades with length. For example, while we can currently obtain around 15,000 CCS reads < 1% from a P6/C4 RS-II run of our 2.6kb *env* amplicon; this read count drops to ~ 1,000 for full-length 9kb HIV genomes.

Both the pipeline and client-side visualizations are under development, with many improvements planned, including a novel clustering algorithm that reduces false positives and a novel consensus algorithm that uses quality scores and performs frame correction. We plan to integrate epitope prediction into the `FLEA` pipeline and add appropriate visualizations for the case when users have IC_50_ values available for their sequences. Finally, `FLEA` will be expanded to support other amplicons.

## Acknowledgments

Research reported here was supported by the National Institute Of Allergy And Infectious Diseases of the National Institutes of Health under Award Numbers R00AI120851 and R01AI120009, and in part by the University of California, San Diego Center for AIDS Research (P30AI036214). K.E. was supported in part by R21AI115701. Computational analysis of sequence data was performed, in part, on a cluster which was supported by U01 GM110749 (NIH/NIGMS). The content is solely the responsibility of the authors and does not necessarily represent the official views of the National Institutes of Health. We are grateful to Sasha Murrell for copy editing.

